# Optogenetic activation of CA1 pyramidal neurons *in vivo* induces hypersynchronous and low voltage fast seizures

**DOI:** 10.1101/2020.09.08.288605

**Authors:** Trong D Huynh, Omar Ashraf, Hayden Craig, Lana Larmeu, Benjemin Barker, Cade Stephenson, Derrick Murcia, Brady Howard, Hai Sun

**Affiliations:** Rutgers Robert Wood Johnson Medical School, Department of Neurosurgery; Louisiana State University Health Science Center, Department of Neurosurgery

## Abstract

Increasing evidence supports the idea that the CA1 of the hippocampus plays an important role in the pathogenesis of temporal lobe epilepsy (TLE). There is however a lack of proof that the over-excitation of CA1 alone is sufficient in inducing seizures *in vivo*. Furthermore, the relevance of the seizures induced from the over-excitation of CA1 to the pathophysiology of TLE is undetermined. Here, we employed optogenetics to activate pyramidal neurons (PNs) in CA1, which reliably induced generalized seizures in freely moving non-epileptic mice. We showed that repeated photostimulations had a kindling effect. In addition, seizures induced by over-active CA1 PNs were dominated by two distinctive onset patterns, i.e. hypersynchronous (HYP) and low voltage fast (LVF) activities, which are widely recorded in patients with and animal models of TLE. In our study, HYP seizures were predominantly associated with the first photostimulation and were entirely replaced by the LVF type afterwards. This phenomenon suggests that the activation of CA1 PNs, when occurring after the first seizure, could lead to the recruitment of GABAergic interneurons to participate in the seizure generation. These findings suggest that seizures induced from the over-excitation of CA1 PNs likely involved the same hippocampal networks and cellular mechanisms underlying TLE.

## INTRODUCTION

Over the past several decades, tremendous progress has been made in understanding the pathophysiology of temporal lobe epilepsy (TLE), which is the most common type of adult onset seizure disorder ^1^. Several mechanisms are implicated in hippocampal excitability:

(1) Neuronal networks serving as the substrate for hypersynchronous seizure activity: There exist three primary excitatory pathways connecting hippocampal and parahippocampal structures. A long tri-synaptic pathway starts from the entorhinal cortex (EC) to the dentate gyrus (DG) to CA3 to CA1, loops back to different layers of EC via subiculum. Two other pathways bypass the DG including an intermediate-length pathway that starts from EC to CA3 to CA1 to subicumlum/EC and a short relay termed temporoammonic pathway that projects directly from EC to CA1 ^2,3^. In normal conditions, most of afferent inputs from EC are filtered and tightly regulated by both the feedforward and feedback inhibitory networks within the DG and CA1 area ^4,5^. As a part of this inhibitory network, both DG and CA1 regions possess GABAergic interneurons to reduce excitatory inputs ^6–11^. Despite powerful inhibitory regulation, these pre-existing excitatory pathways can serve as the substrates for seizure activity as seen in TLE ^12^. The “inhibitory gate” function of DG has been investigated intensively and many have concluded that its dysfunction can allow the generation and propagation of seizure activity resulting in TLE ^3,9,11^. However, the importance of the DG gate theory in TLE has recently been called to question. Several studies have showed that certain animal models of TLE have nearly intact DG ^2,3,6,13^. In addition, the recordings obtained from the brain slices of these animals showed seizures propagated via the temporoammonic pathway to CA1 bypass DG altogether ^2^. Despite the fact that the CA1 region is gaining importance in understanding the pathophysiology of TLE, there is a lack of *in vivo* evidence that the over-excitation of the CA1 pyramidal neuron (PN) alone can lead to seizure.

(2) Different seizure types: Intracranial recordings obtained both from patients with TLE and animal models of TLE have revealed two distinctive seizure onset patterns, defined as low-voltage fast (LVF) and hypersynchronous (HYP) ^14–18^. Furthermore, these two types of seizures have been linked to stereotypical activities of different neuronal subtypes. The LVF seizure, which is more commonly recorded in TLE, is a result of the activity of GABAergic interneurons, whereas, the HYP type is driven by the hyperactivity of glutaminergic neurons including the granule cells at DG and the pyramidal neurons (PN) in the CA1 of hippocampus ^16,18^. These analyses have led to the realization that activities of different neuronal types possess distinct frequency signatures. These analyses, however, have not been carried out in seizures induced by *in vivo* optogenetic manipulation of hippocampal neurons alone. While optogenetics is a powerful tool to investigate the pathophysiology of TLE, its relevance to the actual disease process has yet to be established.

Here, we employed optogenetic techniques to activate the PNs in CA1 and mimic the breakdown of CA1 inhibitory function. We showed that the over-excitation was capable of reliably inducing generalized seizures in free-moving and non-epileptic mice, when the function of DG was entirely unperturbed. We observed a kindling effect of seizure progression after repeated photostimulation. Furthermore, we discovered that both LVF and HYP seizure onset patterns also dominated seizures resulting from activating the PNs in CA1. This finding argued that seizures induced from the over-excitation of the CA1 PNs likely involved the same hippocampal networks and cellular mechanisms underlying TLE. Our study, along with existing evidence, supports the importance of the CA1 region in the pathogenesis of TLE.

## RESULT

### Photostimulation of the CA1 PNs was capable to induce behavioral seizures in non-epileptic mice

To gain optogenetic control over excitatory CA1 PNs, we injected locally an adeno-associated virus AAV5-CaMKIIα-hChR2(H134R)-EYFP (UNC vector core) to deliver the light-activated cation channelrhodopsin (ChR2) under Ca2+/calmodulin-dependent protein kinase II α (CaMKIIα) promotor ^19^ into hippocampal CA1 of healthy wild-type mice (**Fig 1 A,B)**. This method has been proven to express ChR2 mainly on pyramidal/excitatory neurons ^20–22^. Photostimulation (wavelength at 473 nm and 10 Hz pulse) and local field potential (LFP) recording was accomplished by a custom-made hybrid optical fiber with electrode. Behavioral seizures were successfully induced and reproduced by repetitive photostimulation to the CA1 PNs in 7 free moving animals (5 females). All animals were subject to repetitive photostimulation trials. Each trial lasted 90 minutes and comprised three recording segments including 30-minute pre-stimulation observation, 30-minute stimulation, and 30-minute post-stimulation. The 30-minute simulation segment was divided into fifteen 2-minute stimulation epochs. Each stimulation epoch began with a pulsed photostimulation train followed by a resting period **(Fig 1C)**. We estimated the seizure frequency, duration and severity by reviewing the LFP recording, the spectral analysis and the video recording (see methods). The seizure severity score was based on the modified Racine seizure behavioral scores ^12,23–25^.

**Figure 1.**
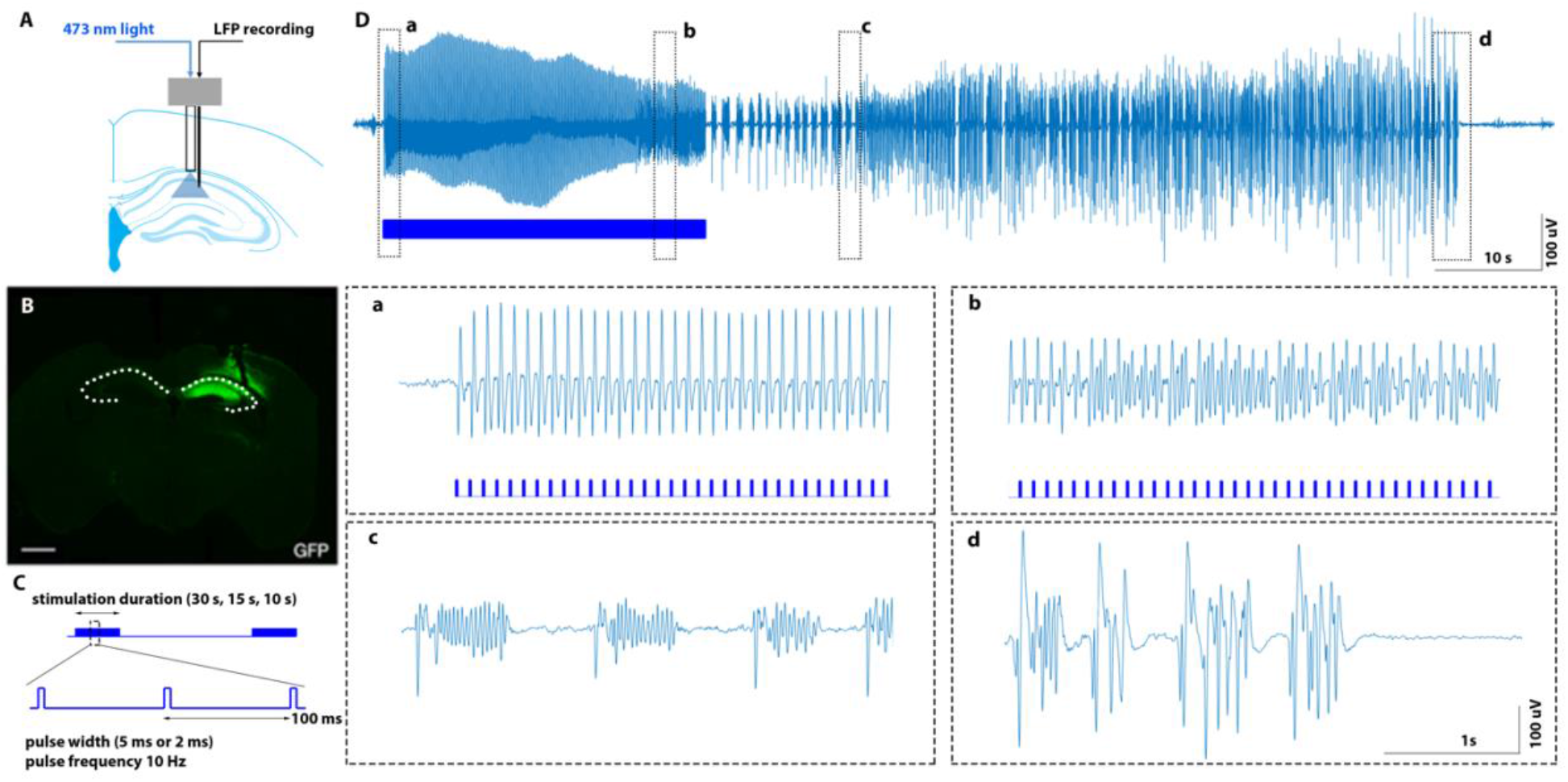
Optogenetic recording setup and characteristic seizure EEG. (**A**) Optrode setup with simultaneous photostimulation and LFP recording of the CA1 region. (**B**) GFP staining showing viral expression restricted in CA1 area of the hippocampus. In addition, the optrode tract is demonstrated with the tip of the optic fiber situated above the CA1 region to ensure illumination of the area (scale bar 1000 μm). (**C)** Photo stimulation scheme with 15 of 2-minute epochs consisting of a 10/15/30 s stimulation period comprised of a 2 or 5 ms light pulse activated at 10Hz. (**D**) A characteristic EEG seizure response recorded with a single electrode in the CA1 area. **(a)** During the stimulation phase, we observed evoked potentials time-locked to each photostimulation pulse (blue bars) without behavioral seizures observed on video recording. Towards the end of the stimulation period (**b**), there was spontaneous EEG activity independent of the photostimulation pulse. After the stimulation period ended, self-sustaining clustered spiking activities arose (**c**), typically associated with motionless behavior (modified racine scale 1-2). These gradually became hypersynchronous discharges (**c**), corresponding to convulsive seizures (modified racine scale ≥ 3) captured on video. The peak hypersynchronous ictal activities were followed by a flattened EEG period which represented the post-ictal state (**d**).

Overall, we successfully induced seizures in 102 out of 109 recording trials (93.6%) among 7 animals. Seizures were detected in 258/1530 (16.86%) stimulation epochs, with the average seizure lasting 46.28 ± 26.09 s. The distribution of the behavioral seizure scores were: 1: 39 (15.12%); 2: 32 (12.40%); 3: 37 (14.43%); 4: 70 (27.13%); 5: 64 (24.81%); 6: 8 (3.10%); 7: 8 (3.10%).

The LFP recorded in the CA1 region showed a stereotypical evolution when a behavioral seizure was induced **(Fig 1D)**. When the photostimulation began, each light pulse resulted in an immediate time-locked evoked potential without behavioral changes observed on video recording, this filed oscillation was defined as optogenetic population discharges (oPD) ^26^ and was observed in all stimulation epochs (**Fig 1Da)**. As the photostimulation continued, the EEG morphology changed with multiple bursts unsynchronyzing to the light pulses (**Fig 1Db)** and continued with self-sustained, high-amplitude, clustered spiking after the photostimulation ended (**Fig 1Dc),** typically associated with motionless behavioral stage (modified Racine scale 1-2). This EEG pattern was defined as optogenetic after-discharge (oAD) ^26–28^. Convulsive seizures took place when the oAD evolved into hypersynchronous discharges. The ictal activities were then followed by depressed EEG wave with lower amplitude compared to baseline, which represented the post-ictal state (**Fig 1Dd)**.

oAD was also induced in some epochs but lasted shortly (< 5s) and did not lead to behavior seizures ^26^ (**Fig 2A)**. No EEG or behavioral changes were recorded when the animals were stimulated with light at a different wavelength (589nm), (**Fig 2C).** Two additional animals were injected with the control virus lacking ChR2 (AAV5-CaMKIIα-EYFP, UNC vector core) and photostimulation in these animals yielded no EEG or behavioral changes (**Fig 2B).**

**Figure 2.**
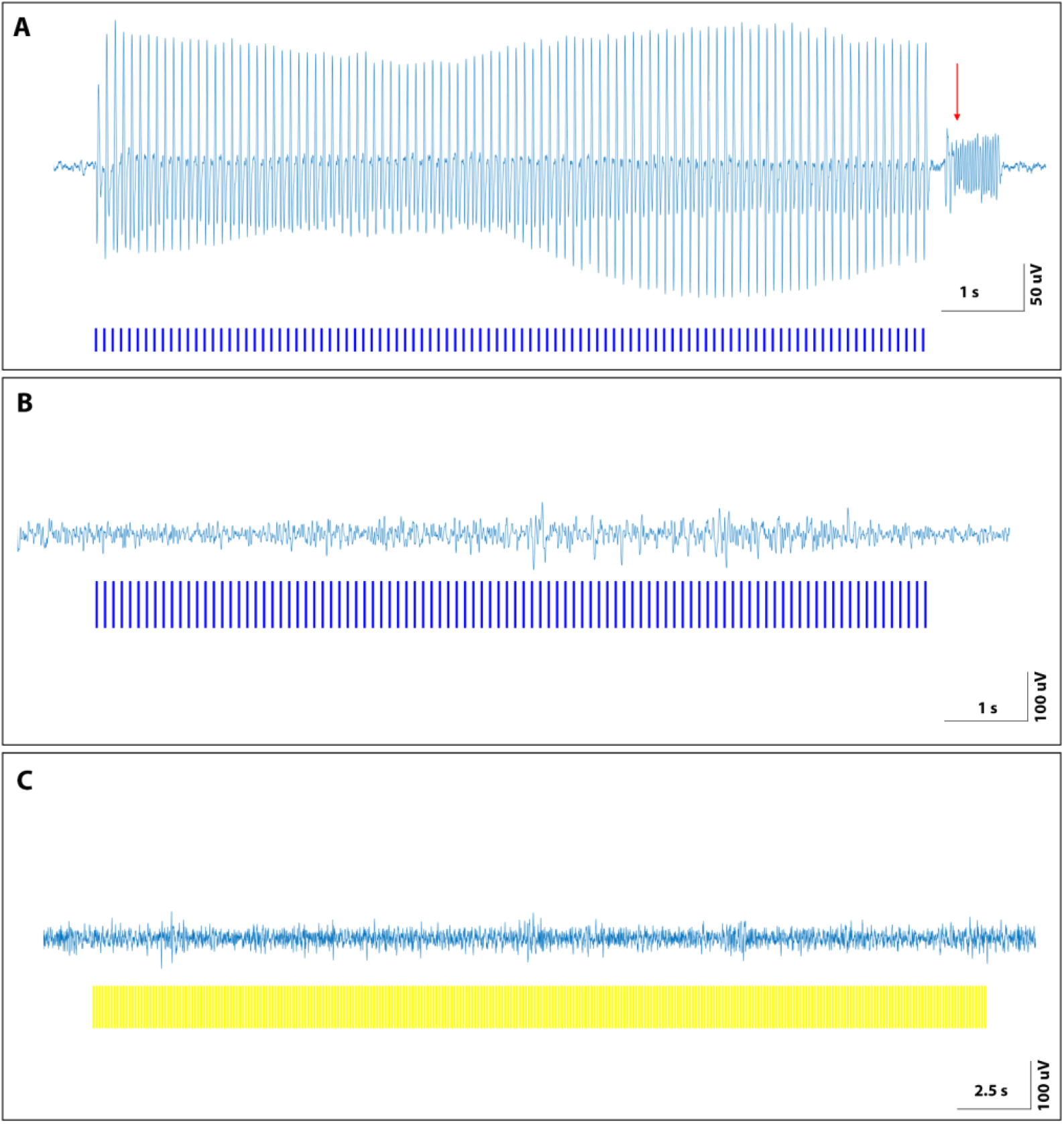
EEG activity in response to photostimulation. **(A)** Shown is an example of an EEG recording in response to blue light photostimulation (473 nm) in a mouse injected with the AAV-ChR2 virus. With each stimulation pulse (blue bar), there is a corresponding evoked potential. There is an afterdischarge (red arrow) following cessation of photostimulation, without associated seizure activity. No EEG changes were elicited in mice with ChR2 expression **(B)** following stimulation with yellow light (589 nm) or in mice with an absence of ChR2 expression **(C)** with blue light photostimulation.

In search for an optimal setting in inducing seizures in these animals, we quantitatively tested the photostimulation with various pulse parameters and train durations. We selected two light pulse width (2ms and 5ms, both at 10Hz) in combination with three light stimulation train durations (10s, 15s and 30s) (**Figure S1**), which generated six permutations of light stimulation settings. Our analyses revealed that the efficacy in inducing seizures did not vary significantly among the trials employing 5ms and 2ms pulse width. The efficacy in inducing seizures was significantly higher with longer stimulation duration. The detailed analyses were included in supplementary materials. Based on these analyses, we selected only the recording trials that had the most effective stimulation settings (10Hz, 30s stimulation duration, 5ms or 2ms pulse) for further analysis. A total of 63 recording trials were available. A total of 181 (19.15%) seizures were detected among 945 recording epochs. The average seizure duration was 49.26 ± 26.12 s. The average seizure severity was 3.56 ± 1.64.

### The kindling effect of the photostimulation of CA1 PNs

The kindling effect of optogenetically induced seizures has been studied by other groups ^12,26,27,29,30^. Here, we examined the kindling effect of accumulative photostimulation of the CA1 PNs by assessing the seizure duration and severity of progression over the time of one recording trial. The same 63 trials included in the previous analysis were included in this analysis. There were a total of 181 seizures among these trials. Each seizure was assigned to the epoch number where the seizures occurred. The seizure duration associated with an epoch was computed by averaging the durations of all seizures occurred in that epoch. The seizure severity associated with an epoch was computed by averaging the severity scores of all seizures occurred in that epoch. The seizure duration increased with repetitive photostimulation (R=0.70, p=0.0048, Spearman’s Rank-Order Correlation) (**Fig 3A, 3B)**. However, the seizure severity did not significantly worsen with repetitive stimulation (R=0.31, p=0.2650, Spearman’s Rank-Order Correlation) (**Fig 3C)**.

**Figure 3.**
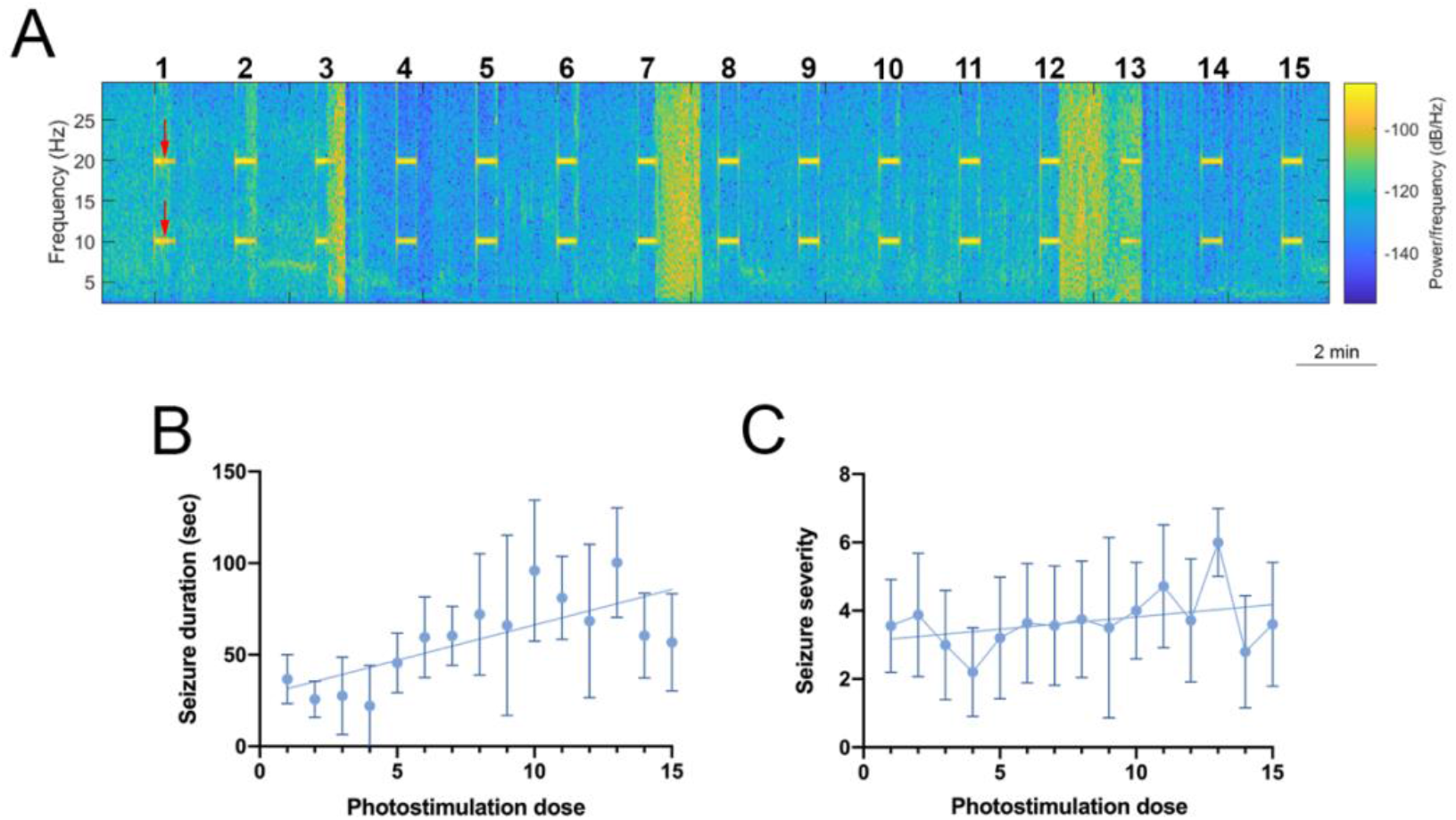
Progression of seizure duration and severity within a trial. The spectrogram demonstrated in **A** shows 5 seizures at epoch 1, 2, 3, 7, and 12 (increased signal power) with progressive lengthening of the seizure duration throughout the recording trial. The stimulation frequency is shown by the red arrows. Seizure duration and severity are quantitatively assessed within a trial in animals that underwent a 30 second stimulation period in **B** and **C**. (**B**) With repeated photostimulation, the seizure duration increased (R=0.70, p=0.0048, Spearman’s Rank-Order Correlation), showing the kindling effect. (**C**) While seizure severity increased with the photostimulation dose, the correlation was not statistically significant. (R=0.31, p=0.2650, Spearman’s Rank-Order Correlation).

### Two onset patterns of seizure induced by optogenetic stimulation

Existing evidence suggested that seizures in patients or animal models of temporal lobe epilepsy (TLE) have two specific onset patterns ^14–18,31–33^. The most frequent onset pattern is characterized by “low-voltage fast” (LVF) activity in the gamma range, at times initiated by a sentinel spike; the second onset pattern, referred to as “hypersynchronous” (HYP), is associated with an initial series of large-amplitude spikes that occurs at a frequency of approximately 1 Hz. Furthermore, studies have suggested that these seizure onset patterns were a result of activities associated with specific neuronal networks ^16,32,34–40^. In this study, we examined the onset patterns of seizures resulting from optogenetically activating the CA1 PNs in healthy mice. Using the previous reported description of two seizure-onset types ^14,17,18^ and examining the EEG morphology and the changes in spectral domain (see methods) in our experiments, we found that optogenetic stimulation of PNs in CA1 also resulted in two distinct seizure-onset patterns recorded in patients with TLE and animal models of TLE, namely HYP and LVF (**Fig 4)**.

**Figure 4.**
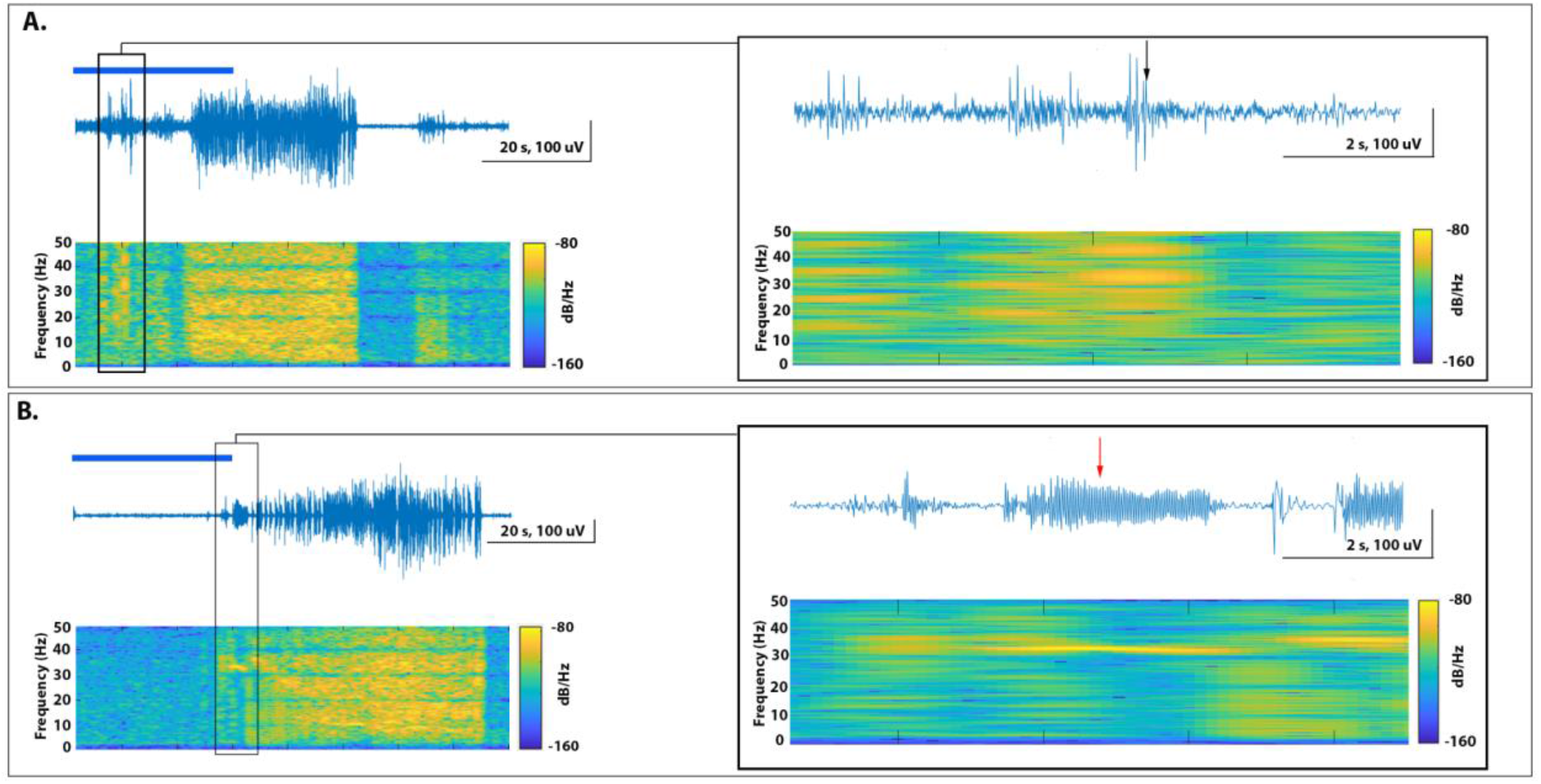
EEG characteristics of HYP and LVF seizure onset patterns. (**A**) HYP seizures were typified by seizure onset early in the stimulation duration (overlying blue bar) with characteristic wide frequency activity noted on the spectrogram. The black arrow on the right shows a high amplitude discharge commonly seen in this seizure type. (**B**) In comparison, LVF seizures were characterized by isolated high frequency activity preceding seizure, usually occurring towards the end of the stimulation duration. The red arrow highlights the high frequency and low amplitude discharge typical of LVF seizures.

As stated previously, we included 181 recorded seizures in this analysis. 63 seizures (34.8%) were HYP, and 97 seizures (53.6%) were LVF, whereas 21 seizures (11.6%) were undetermined. Both seizure types were found within every recording trial. Among 63 HYP seizures, 47 (74.6%) occurred during the 1st stimulation epochs; among 97 LVF seizures, 87 (89.7%) occurred during epochs from 2 to 15. In addition, two seizure types were associated with different timing of the seizure onset relative to the start of photostimulation. HYP seizures on average began 14.76 ± 6.75 seconds after the start of photostimulation, whereas LVF seizures occurred on average 23.29 ± 5.95 seconds (p<0.001, unpaired t test) after the start of photostimulation. Furthermore, two types of seizures had differences in their durations. The average duration for HYP seizures was 34.84 ± 13.37 seconds whereas LVF seizures lasted on average 58.63 ± 32.62 seconds (*p* < 0.001, unpaired t test). The average behavioral scores of two types seizures were, however, not different (type 1 seizure, 3.37 ± 1.35 seconds, type 2 seizure, 3.69 ± 1.80 seconds, *p* = 0.2207, unpaired t test) (**Fig 5**).

**Figure 5.**
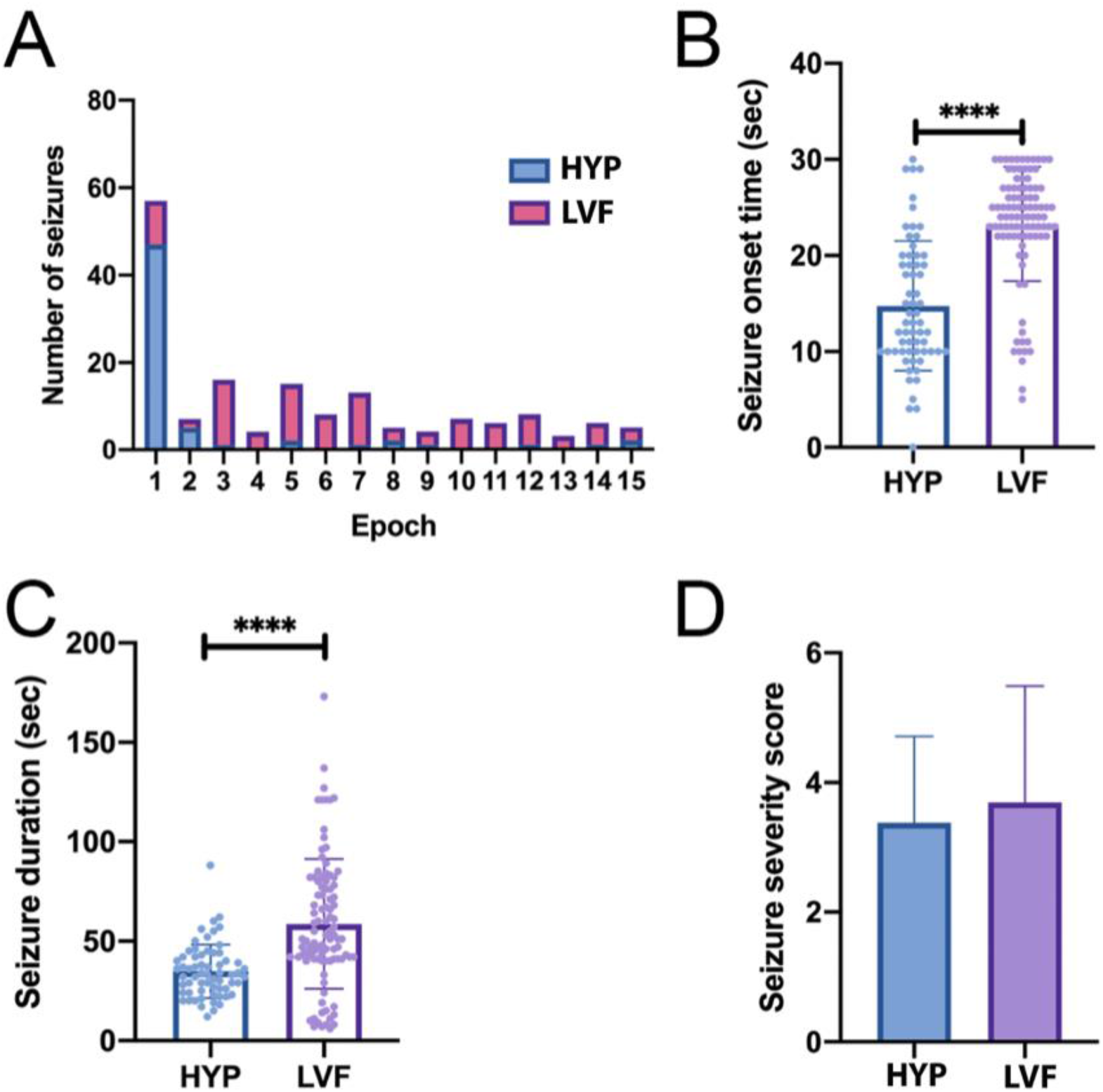
Characterization of HYP and LVF seizures following 30 s stimulation. **(A)** Seizures in epoch 1 were more commonly categorized as HYP seizures, with 82.5% of these seizures exhibiting a HYP pattern. In contrast, 84.5% of seizures observed in epochs 2-15 were characterized by a LVF onset pattern. **(B)** The time from start of photostimulation to seizure onset was significantly shorter in HYP seizures when compared to LVF seizures (14.76 ± 6.75 vs 23.29 ± 5.95 s, respectively; *p* < 0.001, unpaired t test). **(C)** The average seizure duration for HYP seizures was 34.84 ± 13.37 s. This was significantly shorter than LVF seizures, which lasted an average of 58.63 ± 32.62 s (*p* < 0.001, unpaired t test). **(D)** There was no significant difference in the severity between HYP and LVF seizures (3.37 ± 1.35 vs 3.69 ± 1.80, respectively; *p* = 0.2207, unpaired t test).

## DISCUSSION

This study found that (1) photostimulation of channelrhodopsin-expressing PNs in the CA1 region of the hippocampus is capable of inducing seizures in awake, free moving and non-epileptic mice, (2) the photostimulation of CA1 PNs has a weak but definitive kindling effect on epileptogenesis, and (3) seizures resulting from the CA1 PN overactivation have two distinctive seizure onset patterns, hypersychronous (HYP) and low voltage fast (LVF) activities. These results indicate that the CA1 area is a critical region in the hippocampal circuit that plays an important role in seizure genesis and propagation.

### The CA1 region as a gate for controlling excitatory inputs

The hippocampal formation is important for many cognitive functions, such as spatial learning and memory. The hippocampus is also important in pathologies such as temporal lobe epilepsy (TLE). One explanation is based on the idea that the hippocampus contains inhibitory filters or gates preventing too much information from corrupting memory formation as well as preventing seizures ^2,7,41,42^. The dentate gate theory, which emphasized seizure propagation pathway through DG and its important role in controlling seizures, has dominated research filed in epileptic seizures circuitry ^43,44^. Krook-Magnuson and colleagues showed the disruption of gating at DG can lead to seizures and the restoration of the DG ‘gate’ by hyperpolarization of these GCs can stop spontaneous seizure in kainic acid induced TLE in mice ^12^. These experiments supported the DG ‘gate’ theory in TLE.

Here, we showed that the disruption of the gating at CA1 by the over-activation of the pyramidal neurons (PNs) can also lead to seizures in non-epileptic mice. Using optogenetics activating the CA1 PNs to induce seizures *in vivo* has also been reported by another group ^26^. Their findings along with the results presented here support the CA1 being another ‘gate’ among the hippocampal networks. Furthermore, unlike DG, CA1 is involved in all excitatory pathways in these networks: a long trisynaptic loop from entorhinal cortex (EC) to DG to CA3 to CA1 loop back to different layers of EC via subiculum and two additional loops that bypass DG altogether but involves CA1 ^2,3^.

Interestingly, several studies using animal models, showed TLE still occurs among animals with normal DG or the pathological changes in DG associated with TLE have been suppressed ^13^. In addition, *in vitro* studies showed that when the DG is normal, seizures could still spread via the temporoammonic pathway to the CA1 region, bypassing DG altogether ^2^. Conversely, seizures can be effectively controlled among animals with KA-induced TLE when the gating function of the CA1 is enhanced using optogenetic ^45^. Our study, along with existing evidence, seems to suggest that the CA1 region may play a far more important role in controlling excitatory inputs to the hippocampus and in epileptogenesis than previously recognized. The CA1 region may become another therapeutic target for controlling TLE. Examining different targets to control TLE is important since the hippocampus is responsible for important cognitive functions. Having multiple therapeutic targets may allow future treatment strategies that yield better seizure control without sacrificing cognitive functions ^46–48^.

### Optic stimulation associated kindling

Studies including ours have reported kindling effects arising from optogenetically-induced seizures ^12,26,27,29^. In our study, the seizure duration showed a statistically significant increase with repeated stimulation during each experimental recording session. The seizure severity showed a trend of worsening with repeated stimulation; however, this trend did not reach statistical significance. Compared to the electrical kindling, the opto-kindling seems weaker. Existing evidence demonstrated that long-term (24hrs) optogenetic stimulation of CA1 produces changes in spine density and synaptic plasticity ^49^. Recently, an optogenetic kindling model for neocortex epilepsy was also developed and suggested that long term, repeated optogenetic stimulation of the brain produces changes in plasticity and homeostatic regulation of brain networks ^29,50,51^. However, other studies found that the seizure threshold was stable over time ^26,52^. This seems to suggest that the impact of optogenetic activation of hippocampal PNs on epileptogenesis remains to be established.

### HYP and LVF seizures

Our EEG and spectral analyses revealed that there were two distinctive types of seizure onset patterns dominating all seizures resulting from the activation of CA1 PNs, **Fig 5**. These two types of seizures are similar to the two types recorded from patients with TLE and animal models of TLE, i.e., HYP and LVF ^16,36,38,53^ and animal models of TLE ^17,33–35^. To our knowledge, this is the first analysis on the onset patterns for seizure induced by *in vivo* optogenetic manipulation of hippocampal neurons alone. Previously, this type of analysis was carried out in experiments where the optogenetic techniques were combined either with the administration of chemical convulsant such as 4-amiopyridine (4AP) ^18^ or with animal models of TLE ^17^. To our surprise, the onset patterns observed among seizures induced by *solely* activating the CA1 PNs bore close resemblance to the seizures recorded among patients with TLE and animal models of TLE, i.e. HYP and LVF.

Furthermore, the HYP and LVF seizures recorded here have distinctive differences in the timing of the occurrence and the seizure duration. In our study, the HYP seizure was typically seen with the first stimulation epochs whereas the LVF seizures were associated with subsequent stimulation epochs, **Fig 5A**; the HYP seizures were associated with significantly shorter interval between the start of light stimulation and the beginning of seizure activity than the LVF seizures, **Fig 5B**; and the LVF seizures lasted longer than the HYP seizures, **Fig 5C**. Existing evidence suggests that these two types of seizures have been linked to activities of different neuronal subtypes leading to the ictal onset. The LVF seizure is a result of the activity of GABAergic interneurons, whereas, the HYP type is driven by the hyperactivity of glutaminergic neurons ^18^. The fact that the optical stimulation in our experiments directly and precisely activates the glutaminergic neuron, i.e., the PNs, in the CA1 supports the phenomenon that the first seizure is of the HYP type. But the subsequent seizures onset, i.e. the LVF type is a result of the activation of GABAergic interneurons. This phenomenon suggests that the activation of the CA1 PNs, when combined with the first photostimulation-induced seizure, has led to the involvement of GABAergic interneurons in the subsequent seizure generation. In our experiments, the interneurons were not directly activated by the light-activated cation channelrhodopsin (ChR2) under Ca2+/calmodulin-dependent protein kinase II α (CaMKIIα) promotor, which has been demonstrated to have high affinity to glutaminergic neurons ^18^. The activation of interneurons, ushered by the LVF seizure onset pattern, might be a result of the feedforward and feedback inhibition circuits well documented in the CA1 ^17,18,54,55^. The involvement of the interneurons may also be related to the delay in the seizure onset associated with the LVF seizure compared to the HYP seizure, (**Fig 5B)**. While these speculations and hypotheses will need to be studied and confirmed, our data argued that seizures induced from the over-excitation of the CA1 PNs likely involved the same hippocampal networks and cellular mechanisms underlying TLE. Our study could help expand the toolbox of *in vivo* model for TLE, where its pathophysiology can be investigated.

## METHOD

### Animals

Animal use and experimental procedures were carried out according to the protocol approved by the Institutional Animal Care and Use Committee at Louisiana State University Health Sciences Center-Shreveport. A cohort of 9 wild-type Black Swiss (Tac:N:NIHS-BC) mice were used for experiments in this study. The wild-type mice were obtained from crosses of heterozygous *Kcna1*-null animals, which have been described previously ^56^. WT genotypes were confirmed using PCR genotyping of genomic DNA isolated from tail clips, as done previously ^56^. Mice were housed in controlled conditions (20-25 °C, 12 h light and dark cycle with free access to food and water).

### Optrode assembly for simultaneously optogenetic stimulation and local field potential recording

The optrode assembly was a custom-made design, using a 3-channel electrode (PlasticsOne; VA, USA) and an optic cannula Zirconia ferrule 1.25 mm, fiber optic 200 μm core-diameter, 0.37 NA (Doric Lenses; Quebec, Canada). The electrode pedestal and optic cannula were held together by a custom-made 3-D printed sleeve to ensure consistent placement and alignment. One of the channels was aligned alongside the optic fiber to achieve photostimulation and local field potential (LFP) recording simultaneously. Of note, the tip of the electrode was extended 0.5 mm beyond the tip of the optic fiber to ensure the illumination and recording in a same area.

### Stereotaxis viral infection and chronic optrode implantation

This study employed two-staged surgeries: the first surgery for viral injection and the second surgery for optrode implantation. Mice (n = 9) were anesthetized by isoflurane (2.5%, 97.5% oxygen, 0.5 L/min), administered subcutaneously buprenorphine (1 mg/kg) followed by carprofen (1 mg/kg) to minimize pain, then positioned into the stereotaxic frame (Model 942, David Kopf Instruments, Tujunga, CA, USA). A 3-cm midline skin incision was made to expose the skull landmarks. A burr hole was created over the CA1 subfield of the hippocampus (anteroposterior, AP: −1.9 mm; mediolateral, ML: +1.6 mm from bregma) (Franklin & Paxinos 2007) to allow for viral injection. For the experimental group (7 mice), AAV5-CaMKIIɑ-hChR2(H134R)-EYFP (1×10^13^ vg/mL; UNC vector core) was infused into the hippocampal CA1 (dorsoventral DV – 1.6 mm from dura) to express Channelrhodopsin2-EYFP (ChR2-EYFP) fused protein primarily in excitatory neurons ^20–22^ under control of the Ca2+/calmodulin-dependent protein kinase IIɑ (CaMKIIɑ) promoter. For the control group (2 mice), control virus AAV5-CamKIIɑ-EYFP (titer 1×10^13^ vg/mL; UNC vector core) was injected with identical brain coordinates. Following injection of 2.5 μl virus suspension by a 34-gauge needle and Hamilton syringe, the needle was kept in place for 5 minutes to avoid back-flow. The burr hole was covered with bone wax, and the incision was closed using veterinary-grade cyanoacrylate glue. The animal was then allowed to recover from the procedure and was then monitored while the viral expression took place.

After an interval of approximately 3 weeks (mean 3.4 +/−0.9 weeks), the animal underwent the optrode implantation. The optic fiber attached electrode wire was lowered into the brain near the previously viral injection site (AP −2.0 mm; ML +1.6 mm). The electrode tip and optic fiber tip were situated at 1.6 mm and 1.1 mm from the dura, respectively (**Figure 1A**). Two additional burr holes were also drilled to insert the reference electrode on the cortical surface (AP −4.0 mm, ML +1.6 mm) and ground screw (AP −4.0 mm, ML −1.6 mm). Once all components were in place, dental cement was used to secure the assembly on the skull. The skin incision was closed around the assembly using veterinary-grade cyanoacrylate glue. The animals were then allowed to recover for at least 7 days from the second surgery before the experiment of photostimulation and electrophysiology recording.

### Photostimulation and local field potential recording

Simultaneous photostimulation and LFP recording was performed. Recordings took place in a standard plastic “shoebox” cage with the lid removed. Mice were tethered via a fiber optic cable and electrode cable. ChR2 excitation was performed using a 473 nm laser (LaserGlow Technologies, Otanrio, Canada) with a terminal output of approximately 5-6 mW. Each trial lasted 90 minutes and comprised three recording segments including a 30-minutue pre-stimulation observation, a 30-minute stimulation, and a 30-minute post-stimulation. The 30-minute simulation segment was divided into fifteen 2-minute stimulation epochs. Each stimulation epoch began with a pulsed photostimulation train followed by a resting period. Experimental parameters included stimulation train duration and pulse width. Two light pulse widths (2ms and 5ms, both at 10Hz) in combination with three light stimulation train durations (10s, 15s and 30s) (**Figure 1C**), which generated six permutations of light stimulation settings namely setting 1: 30s-5ms; setting 2: 15s-5m; setting 3: 10s-5ms; setting 4: 30s-2ms; setting 5 15s-2ms; setting 6: 10s-2ms. Each animal was also tested with non-corresponding amber light (589 nm, 10Hz, 5 ms pulse, 30s train). Signals were amplified and sampled at 3 kHz, low-pass filtered at 500 Hz. Photostimulation parameters control and LFP recordings were performed using RZ5D processor and Synapse software (Tucker-Davis Technologies, Florida, USA).

### Seizure analysis

Field potential signals were band-pass filtered in 0 to 100 Hz frequency range and notch filtered at 60 Hz. Time-frequency analyses were performed by spectrogram using short-time Fourier transform (Signal Processing Toolbox Released 2019b, MATLAB 9.7.0, MathWorks, MA, USA). All electrography and video recordings were reviewed manually by three independent reviewers (HS, TDH and OA). Seizures were determined on raw LFP signal plot and spectrogram as the appearance of afterdischarges (asynchronous spiking, increasing amplitude, frequency, and signal energy compare to baseline) lasting more than 5 seconds. Severity of behavioral symptoms for each electrographic seizure was scored using synchronized video according to modified Racine scale ^12,23–25^: stage 1: a change in behavioral state (sudden behavioral arrest or sudden motion); stage 2: head nodding; stage 3: forelimb clonus; stage 4: rearing, or clonus when on belly, or strong hindlimb clonus (bucking); stage 5: falling, or clonus when on side; stage 6: multiple sequences of rearing and falling, or brief jumps; stage 7: violent jumping; and stage 8: class seven, followed by a period of tonus lasting longer than 5 seconds.

For determination of seizure-onset patterns, notch filter was applied at 10Hz to truncate the appearance of field oscillation i.e. population discharges at a similar frequency of the pulsed photostimulation. Seizures were classified into 2 groups according to their onset patterns that reported by previous studies ^14,17,18,33^, defined as low-voltage, fast (LVF) and hypersynchronous (HYP). LVF seizures are characterized at onset by the occurrence of a sentinel spike followed by low-amplitude, high-frequency activity, whereas the initiation of HYP seizures coincides with a series of focal (so-called pre-ictal) spiking at a frequency of approximately 2Hz.

### Immunohistochemistry

Following optogenetic stimulation, subjects were lethally anesthetized using isoflurane (5%). The subjects were then perfused transcardially with 0.01M phosphate-buffered saline (PBS) followed by 4% phosphate-buffered paraformaldehyde. The brain was further fixed for 24 hours at 4°C in 4% paraformaldehyde. The sample was transferred to 30% phosphate-buffered sucrose for cryoprotection for 36 hours and then embedded using POLARSTAT PLUSTM Frozen Embedding Medium at −20°C. A cryostat (Leica CM3050 S; Leica Biosystems Inc., Buffalo Grove, IL, USA) was used to slice the brain into coronal sections (50 μm thickness) and stored in a 24-well plate containing 1 mL of 0.01M PBS at 4°C until staining. The Mouse Brain in Stereotaxic Coordinates (4th edition) was referenced for selecting representative tissue sections of the hippocampus. 4’, 6-diamidinio-2-phenylindole (DAPI; ThermoFisher Scientific, Little Rock, AR, USA) was administered to the sections to label neuron body. Sections were then mounted onto gelatin subbed slides using Dako Fluorescent Mounting Medium and cover-slipped. The samples were visualized with a microscope (Zeiss AxioObserver with Apotome; ZEISS, Oberkochen, Germany) to qualitatively confirm viral expression and optrode placement.

### Data analysis and statistics

Data was presented as mean ± standard deviation (SD) unless stated otherwise. Statistical analysis was performed in Prism 8 (version 8.3.1, GraphPad Software). Unpaired t-test was used to compare differences between two groups. One-way ANOVA test was used to examine the difference between 3 or more groups. A correlation between the number of repeated photo-stimulation and seizure duration or severity was tested using Spearman’s correlation coefficient. P < 0.05 was considered statistically significant.

## Acknowledgements

This work was supported by an Idea Development Award in Epilepsy Research Program from the Department of Defense Congressionally Directed Medical Research Programs (CDMRP), W81XWH-18-1-0655.

## Author contribution

H.C, L.L, B.B, C.S, D.M, B.H: carry out in vivo and immunohistochemistry experiments, collected data; O.A: analyzed data, prepared figures; T.D.H: Conceptualized study, analyzed data, prepared figures, wrote the manuscript; HS: Supervised, conceptualized and designed the experiments, wroted and final approve the manuscript. All authors reviewed the manuscript.

## Supplementary materials

In search for an optimal setting in inducing seizures in these animals, we quantitatively tested the photostimulation with various pulse parameters and train durations. We selected two light pulse width (2ms and 5ms, both at 10Hz) in combination with three light stimulation train durations (10s, 15s and 30s), which generated six permutations of light stimulation settings. The parameters for each stimulation setting were the following (stimulation duration - pulse width): setting 1: 30s-5ms; setting 2: 15s-5m; setting 3: 10s-5ms; setting 4: 30s-2ms; setting 5 15s-2ms; setting 6: 10s-2ms. The probability in eliciting a seizure in each recording session was as follows: setting 1: 100% (54/54); setting 2: 88.2% (15/17); setting 3: 81.8% (9/11); setting 4: 100% (9/9); setting 5: 100% (8/8); setting 6: 70% (7/10). Photostimulation with 30s-5ms and 30s-2ms resulted in 2.79 ±1.37 (n = 54) and 2.89 ± 1.45 (n = 9) seizures per recording session, respectively. Photostimulation with 10s-2ms had the lowest number of seizures per session, 1.00 ± 0.94 (n = 10). Our results showed seizure frequency was dependent on light stimulation settings (p=0.0009, One-way ANOVA; **Fig. S1A**) and 30s-5ms and 30s-2ms are the two most effective stimulation settings. Similarly, seizures duration was longest when induced by 30s-5ms setting (50.21± 28.15, n = 155), and shortest when induced by 10s-5ms or 10s-2ms, (34.88 ± 22.70, n = 17; 33.50 ± 13.90, n = 10). Seizures duration was also dependent on stimulation settings (p=0.0163, One-way ANOVA; **Fig. S1G**). Seizure severity behavioral score, however, was not statistically different among stimulation settings (p=0.8586, one-way ANOVA; **Fig. S1D**).

We grouped the recording sessions by light pulse either 5 ms (n = 82) or 2 ms (n = 27). Seizure frequency in 5 ms pulse is not significantly different to that in 2 ms pulse, 2.49 ±1.48 (n = 82) vs 1.856±1.406 (n = 27) (p=0.0521, unpaired t test; **Fig. S1B**). Average seizure duration was 48.47 ± 27.35 (n = 208) and 37.17 ± 17.76 (n = 50) in 5 ms pulse and 2 ms pulse, respectively (p=0.0050, unpaired t test; **Fig. S1H**). For seizure severity, the average behavioral score of seizures induced by 5 ms pulse and 2 ms pulse was 3.71 ± 1.36 (n =208) and 3.96 ±1.54 (n = 50), respectively (p=0.6574, unpaired t test; **Fig S1E**). Our data showed that while pulse width affected seizure duration, it had no effect on seizure frequency and seizure severity.

We also evaluated seizure metrics regarding stimulation length. The recordings were then grouped by 30s length (n = 63), 15s length (n = 25), and 10s length (n = 21). Seizure frequency was highest in 30s length (2.81 ± 1.37, n = 63) and lowest in 10s length (1.29 ± 1.31, n = 21). Seizure frequency was dependent on stimulation length (p<0.0001, one-way ANOVA; **Fig. S1C**). Average seizure duration induced by 30s, 15s, and 10s photostimulation length were 48.77 ± 27.02 (n = 181), 43.46 ± 24.02 (n = 52), 34.37 ± 19.60 (n = 27), respectively. Seizure duration was also dependent on stimulation length (p=0.0194; one-way ANOVA; **Fig. S1I**). Average behavioral score was: 3.55 ± 1.60 (n = 181); 3.87 ± 1.24 (n = 52); and 3.58 ± 1.24 (n = 27); in 30s, 15s, and 10s, respectively. In contrast with seizure frequency and seizure duration, seizure severity was not dependent on stimulation length (p=0.4234, one-way ANOVA; **Fig. S1F**).

---

**Figure 1S.**
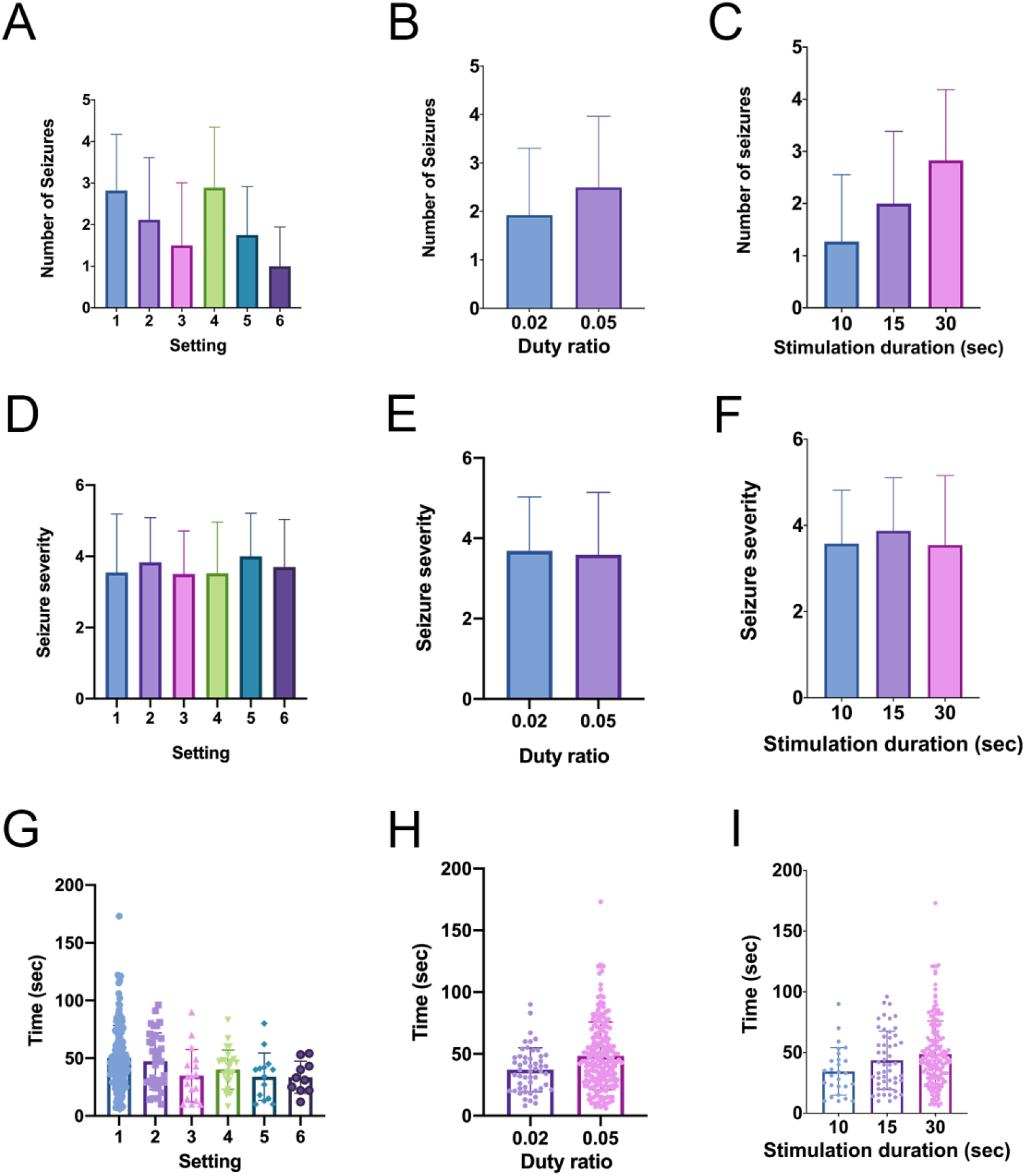
Influence of stimulation setting, duty ratio, and photostimulation duration on seizure number, severity, and duration. The average number of seizures per trial were quantified in **A**, **B**, and **C**. (**A**) Analysis revealed that seizure number was dependent on the photostimulation settings (p=0.0004, One way ANOVA). (**B**) Assessing seizure number by the duty ratio demonstrated that it has no effect on the seizure number (p=0.0816, unpaired t test). In contrast, stimulation duration was associated with a greater number of seizures per trial (p<0.0001, One way ANOVA). Seizure severity was evaluated in **D**, **E**, and **F**.(**D**) The photostimulation settings had no effect on average seizure severity (p=0.8449, One way ANOVA). Similarly, seizure severity was not dependent on duty ratio (**E**; p=0.704, unpaired t test) or stimulation duration (**F**; p=0.4025, One way ANOVA). Seizure duration was assessed in **G**, **H**, and **I**. (**G**) Seizure duration was dependent on the photostimulation setting (p=0.0190, One way ANOVA). (**H**) A duty ratio of 0.05 resulted in a longer seizure duration than a duty ratio of 0.02 (p=0.0066, unpaired t test). (**I**) Also, there was a statistical difference in seizure duration when grouping seizures by photostimulation duration (p=0.0194; One way ANOVA).

